# Characterization of popcorn temperate and tropical populations and GWAS for zeins and starch contents

**DOI:** 10.1101/2023.03.16.533023

**Authors:** Leonardo Fioravante Gotardi, José Marcelo Soriano Viana, Matheus Pereira Ribeiro, Raissa Barbosa de Castro, Humberto Josué de Oliveira Ramos, Juliana Lopes Rangel Fietto

## Abstract

Because measuring expansion volume (EV) is simple and inexpensive, popcorn breeders have developed high-quality single crosses ignoring the contents of zeins, starch, lipids, and cellular wall components in selection. However, some methods of quantification of these quality-related traits can be applied to popcorn breeding, increasing the selection efficacy for quality. The objectives of this study were assess methods of quantification of zeins and starch that can be used in popcorn breeding, characterize a temperate and a tropical populations for zeins and starch contents, and identify candidate genes for these quality-related traits. We genotyped and phenotyped 286 plants. For quantification of total zeins and zein subunits we choose the ‘lab-on-a-chip’ microfluidic electrophoresis. For quantification of starch and amylose/amylopectin we choose the Megazyme’s Amylose/Amylopectin kit assay. The temperate population has superior EV (+36%), a higher level (+32%) of the 19 kDa zein subunit, and lower levels of the 21, 22, and 27 kDa subunits (−1543, −40 and −47%, respectively). Although there are statistical differences between the two populations regarding starch, amylose, and amylose/amylopectin ratio, the differences are not significant (−2 to 8%). Six candidate genes were identified for the 19 and 22 kDa zeins, one for the 21 kDa zein, one for total zeins, two for starch, and four for amylose, with emphasis on three genes from the Z1C subfamily, coding for the 19 and 22 kDa alpha-zeins, located on chromosome 4. The evaluated quantification methods can be used in popcorn breeding programs, with the potential to revolutionize the breeding for quality.

**Key message:** The protein chip and the Amylose/Amylopectin kit assay for zein and starch quantification, respectively, can be effectively used in popcorn breeding, with the potential to revolutionize the breeding for quality.

## Introduction

Maize grains are composed of starch (72%), proteins (10%), lipids (4%), vitamins, and minerals (Nuss and Tanumihardjo 2010). The expansion volume (EV; mL/g) and aspects related to palatability, such as texture and flavor, are the major quality traits of popcorn kernels (Sweley et al. 2013). Starch is an essential polymer involved in the grain’s expansion process (Sweley et al. 2013) and consists of the homopolysaccharides amylose and amylopectin (Oliver and Pan 1995). Amylose accounts for about 20 to 30% of the starch content in maize grains and is characterized as a linear glucose chain with predominantly α-1,4 bonds, with occasional branches due to α-1,6 bonds. Amylopectin is a more complex glucose polymer due to the predominance of branches in the chain with α-1,6 bonds between glucose residues and represents about 70 to 80% of the starch content of the grain (Oliver and Pan 1995). Although there is no significant correlation between EV and amylose, amylopectin (Borras et al. 2006) and starch (Saito et al. 2021) contents, these compounds are associated with grain texture and have a critical function during the expansion of the translucent endosperm.

The maize grain storage proteins are classified into four families by primary sequence similarity, denominated albumins, globulins, glutelins, and prolamins. The prolamins are the principal and most abundant protein in the endosperm (50 to 70%) (Li and Song 2020) and are characterized by having a higher proportion of proline and glutamine. Maize prolamins were characterized and named zeins by John Gorham in 1821. Zeins could be classified into four subunits (α-, β-, γ- and δ-zeins) based on solubility, ability to form disulfide bridges, and molecular mass (Holding 2014). The alpha-zeins (19 and 22 kDa) are encoded by four gene subfamilies (Z1A, Z1B, Z1C, and Z1D) and account for approximately 70% of the total endosperm’s zeins (Song et al. 2001; Song and Messing 2002). Beta-zein (15 kDa), delta-zeins (10 and 18 kDa) and gamma-zeins (16, 27 and 50 kDa) are encoded by a single gene (Li and Song 2020). Gamma-zeins represent the second largest fraction of zeins in the endosperm and contain a high proportion of cysteine. In popcorn, alpha-zeins are related to endosperm hardness and EV, as demonstrated by Borras et al. (2006).

Genome-wide association studies (GWAS) identify genes related to complex traits by detecting significant phenotypic differences between classes of single nucleotide polymorphisms (SNPs). Genome-wide association study represents an advanced and effective method for understanding the genetic architecture of quantitative traits. Zheng et al. (2021) conducted a GWAS in a maize panel with 248 inbred lines and identified 29 candidate genes related to grain quality. Among these genes, three are related to protein content, located on chromosomes 1 and 2, and five are related to starch content, located on chromosomes 1, 3, 4, and 6. Li et al. (2018b), using a panel of 464 maize inbred lines, identified 39 candidate genes related to amylose content, located on chromosomes 3, 4, 5, 8, and 9. Pang et al. (2019) combined genomic and transcriptomic analyses of a panel with 282 maize inbred lines, aiming to understand the molecular basis of grain development. Among the candidate genes, 10 are zein coding and two are known regulators of zein genes or genes determining endosperm development (Opaque2 and ZmICE1).

In popcorn breeding, there is limited information about the contents of zeins, starch, lipids, and cell wall components in grains. The absence of studies involving the quantification of these components on a scale compatible with breeding programs – tens to thousands of individuals or inbred/doubled-haploid lines – should be due to the difficulty of developing quantification methods that are compatible with large-scale evaluations, precise, and economically viable. Thus, the initial aim of this study was to evaluate the ‘lab-on-a-chip’ microfluidic gel electrophoresis as a tool for the quantification of total zeins and zein subunits in breeding. This technique is accurate and compatible with the requirements of breeding programs (Goetz et al. 2004). Another objective was to evaluate the large-scale use of the commercial Amylose/Amylopectin kit from Megazyme (Megazyme Ireland International, Ltd, Bray, Ireland), following the standard protocol. Because the procedure is an enzymatic method, it is precise, reliable and economical.

As the quantification techniques evaluated were used in two populations, one temperate and the other tropical, another aim of this study was to provide a better understanding of the difference in quality between temperate (higher quality) and tropical (lower quality) germplasms, widely recognized by popcorn breeders. Finally, we used genotyping by sequencing (GBS) data to identify and validate genes related to the biosynthesis and accumulation of zeins, starch, amylose, and amylopectin.

## Materials and methods

### Populations

In this study, 183 plants from the temperate population UFV-MP5 and 103 from the tropical population Beija-Flor c4 were genotyped and phenotyped. UFV-MP5 is a biparental population derived by random cross from the North American hybrid AP4502, developed by the Agricultural Alumni Seed Improvement Association, Romney, IN, USA. Beija-Flor c4 was derived after four half-sib selection cycles for EV at the Federal University of Viçosa, Brazil.

### Extraction and quantification of total zein and zein subunits

As the highest concentration of zeins is observed in the endosperm, the extraction protocols involve the removal of the pericarp and embryo from the grain. Seven grams of grains were immersed in distilled water for 30 min to facilitate the removal of the pericarp and embryo. The pericarp was removed with forceps and the embryo was removed with a scalpel. The endosperms were dried in a freeze dryer for 96 h (Landry et al. 2004), followed by grinding in a ball mill. Before the extractions, lipids were removed from the endosperm. This step consisted in adding petroleum ether (1 mL) to the endosperm samples (100 mg), followed by shaking (15 min, 25 °C, and 130 rpm), centrifugation (15 min, 25 °C, and 12000 x g), and removal of the supernatant containing the lipid fraction (Landry et al. 2000). This procedure was performed twice. The defatted flour was then dried in a freeze dryer for 12 h.

Zein extraction was performed by two successive steps: extraction of total protein and extraction of total zeins. The extraction of total proteins was performed by adding 1 mL of extraction buffer (0.0125 mM sodium borate - pH 10; 1% SDS (v/v); and 2% 2-β-mercaptoethanol) to the defatted flour. The microtubes were shaken (25 °C and 130 rpm) for three hours, followed by centrifugation (15 min, 25 °C, and 13,300 x g) and storage of the supernatant (Parsons et al. 2020; Wallace et al. 1990). The supernatant contained the protein portion of the extract and was the fraction used for extracting total zeins. Zein extraction followed the procedure proposed by Wallace et al. (1990), with adjustments for popcorn. The methodology involves the separation of total protein extracts into zein and non-zein fractions (Hamaker et al. 1995; Wallace et al. 1990).

Zeins are soluble in alcoholic solution. The most common extractants are ethanol and isopropanol. Total zeins were extracted by adding absolute ethanol to the supernatant of the total protein extraction to give an extract containing 70% alcohol (v/v) (Wallace et al. 1990). The microtubes were incubated overnight in shaking (16 h, 4 °C, and 30 rpm), followed by centrifugation (15 min, 4 °C, and 13,300 x g), and the supernatant was stored. The supernatant contained the zein portion and the pellet contained non-zein proteins (Parsons et al. 2020; Wallace et al. 1990). The zein extracts were characterized by ‘lab-on-a-chip’ microfluidic electrophoresis. Quantification of total zeins and zein subunits was performed using the Agilent 2100 Bioanalyzer system and an Agilent Protein 80 kit (recommended for proteins between 5 and 80 kDa) according to the Agilent Technologies protocol. Percentage estimates for zein subunits and total zein concentration were determined using Agilent 2100 expert software. Quantifications were based in three biological replicates.

A sample of the plants was also characterized by sodium dodecyl sulfate-polyacrylamide gel electrophoresis (SDS-PAGE), using a commercial zein standard (Sigma-Aldrich) and a molecular mass marker to verify the zein subunit distribution profiles (Parsons et al. 2020). Electrophoresis was performed according to Laemmli (1970), using stacking and separating gels containing 4 and 12.5% polyacrylamide, respectively. After electrophoresis (2 h, 100 V), the gels were fixed in a fixing solution (10% methanol and 5% acetic acid) with shaking (2 h, 25 °C, and 40 rpm). Gels were stained in a Coomassie G-250 solution for 48 h and then added to a decolorizing solution of 7.5% acetic acid (v/v) and 25% methanol (v/v). The gels were scanned to assess the quality of the extractions (Online Resource Figure 1).

**Figure 1.**
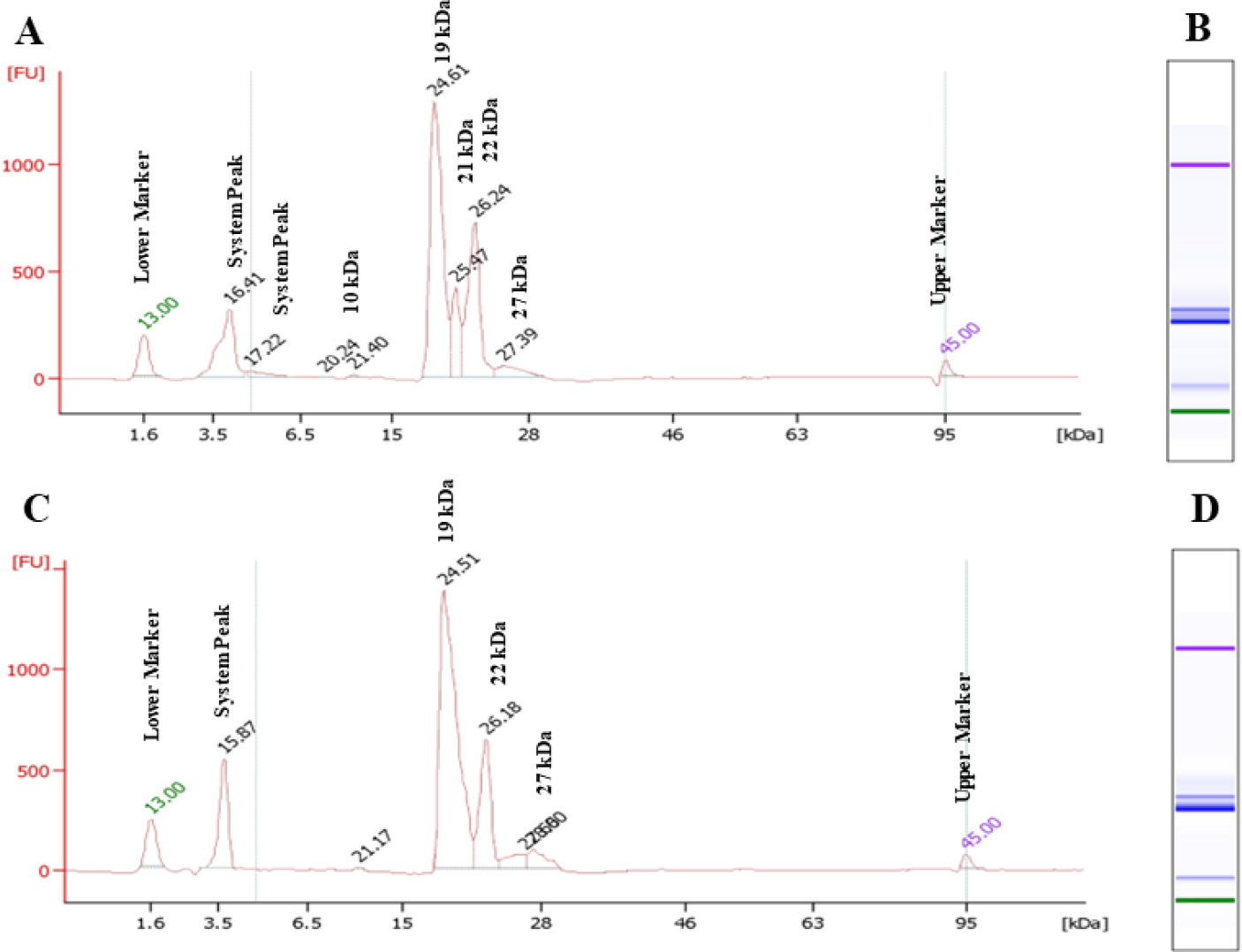
Comparative analysis of zein subunits in the tropical and temperate populations using capillary electrophoresis. A. Electropherogam frequently observed in zein extracts in the tropical population. B. Zein profile of an individual from the tropical population. C. Electropherogam frequently observed in zein extracts in the temperate population. D. Zein profile of an individual from the temperate population.

### Extraction and quantification of amylose, amylopectin, and starch

Extractions were performed with grains previously stored in a refrigerated chamber. Two grams of grains were ground in a ball mill until a homogeneous flour was obtained, followed by extraction with the Megazyme commercial Amylose/Amylopectin Kit (Megazyme Ireland International, Ltd, Bray, Ireland), according to the standard protocol (https://www.megazyme.com/amylose-amylopectin-assay-kit). Absorbance measurements were performed in a microplate spectrophotometer at a wavelength of 510 nm. Total starch content was estimated as a percentage using the formula established by Gibson et al. (1997), which includes standard glucose absorbance. Quantifications were based on two biological replicates and five technical replicates.

### EV quantification

For characterizing the populations in regard to EV, we used the data from Viana et al. (2022) and by Paes et al. (2016). The EV was measured using samples of 10 g and a 27-L microwave oven (900 W).

### Genotyping

Leaf samples for DNA extraction were collected at the vegetative stage 3 (three fully expanded leaves). Genomic DNA was extracted using the CTAB (cetyl trimethylammonium bromide) protocol with modifications (Doyle and Doyle 1990; Jiao et al. 2017). Purity was determined using a NanoDrop spectrophotometer (Thermo Scientific, USA) and DNA quantification was performed using a Qubit® 2.0 fluorometer (Life Technologies). The GBS of the temperate population was performed by the Institute of Biotechnology at Cornell University using a Hi-Seq 2500 sequencer (125 bp paired-end reads). The GBS of the tropical population was performed by the Institute for Research in Immunology and Cancer (IRIC), University of Montreal, using a NextSeq500 sequencer (85 bp single-end reads).

Two SNPs calls were made, one by the Institute of Biotechnology (temperate population) and the other by Omega Bioservices, Norcross, GA (tropical population) using the reference genome B73 version 4 (Jiao et al. 2017). The raw dataset for each SNP call was read using the vcfR package (Knaus and Grunwald 2017), followed by filtering for missing alleles and chromosomes. Then, SNP call rate, genotype call rate, and minor allele frequency (maf) were calculated. Then, we filtered again for maf > 0.01 and call rates > 0.85 and performed the Hardy-Weinberg equilibrium test with Bonferroni correction, followed by imputation with Beagle (Browning and Browning 2009), using the R package Synbreed (Wimmer et al. 2012). After quality control and imputation, the numbers of SNPs in the temperate and tropical population datasets were 145,420 and 109,999, respectively.

### GWAS

We fitted the model proposed by Yu et al. (2006). Population structure analysis was performed using Structure software (Pritchard et al. 2000), using 1000 randomly SNPs (100 per chromosome). The parameters adopted were as follows: the length of burn-in period, 1000; the number of Markov Chain Monte Carlo replications after burn-in, 9000; no admixture model with independent allele frequencies; the number of suggested populations (K), 1 to 4; and five iterations. The number of subpopulations was defined based on Evanno et al. (2005). A subpopulation structure was observed only for the temperate population, with two subpopulations. A total of 138 (75.4%) plants were classified into a subpopulation (Online Resource Figure 2).

The R package rrBlup was used for the association analyses (Endelman 2011). In the analysis of the starch and amylose contents of the temperate population, due to a high number of significant associations identified using rrBlup, the R package GWASpoly was used (Rosyara et al. 2016). To control the type I error, the false discovery rate (FDR) proposed by Benjamini and Hochberg (1995), was used, adopting 5% as the significance level. Probability values were determined using the R package q-value (Debney and Storey 2011).

For candidate genes search, a 120-kb interval upstream and downstream of each significant SNP was used. The functional annotation of candidate genes was performed using the MaizeGDB database (https://www.maizegdb.org/), NCBI (https://www.ncbi.nlm.nih.gov/), and UniProt (https://www.uniprot.org/), with the B73 version 4 genome as a reference. The GO analysis of candidate genes was performed using tools from the Gene Ontology website (http://geneontology.org/).

## Results

Microfluidic electrophoresis allowed the identification of five zein subunits based on their molecular weight (10, 19, 21, 22, and 27 kDa) (Figure 1). In the two populations, these subunits corresponded, on average, to 98% of the bands mapped in the zein extracts. The population means were significantly different concerning the contents of the 19, 21, 22, and 27 kDa subunits. The temperate population has lower concentration of the 21, 22, and 27 kDa subunits (−1543, −40, and −47%, respectively) and a higher content (+32%) of the 19 kDa subunit (Table 1). The total zeins content in the temperate population was 41% higher than that in the tropical population. Although there were significant differences between the two populations regarding the starch, amylose, and amylopectin contents and the amylose/amylopectin ratio, they were not important (−2 to 8%). It is likely, therefore, that the statistical superiority of the temperate population for EV (26%) is due in part to the differences in total zein content and the contents of the 19 to 27 kDa zein subunits.

**Table 1.**
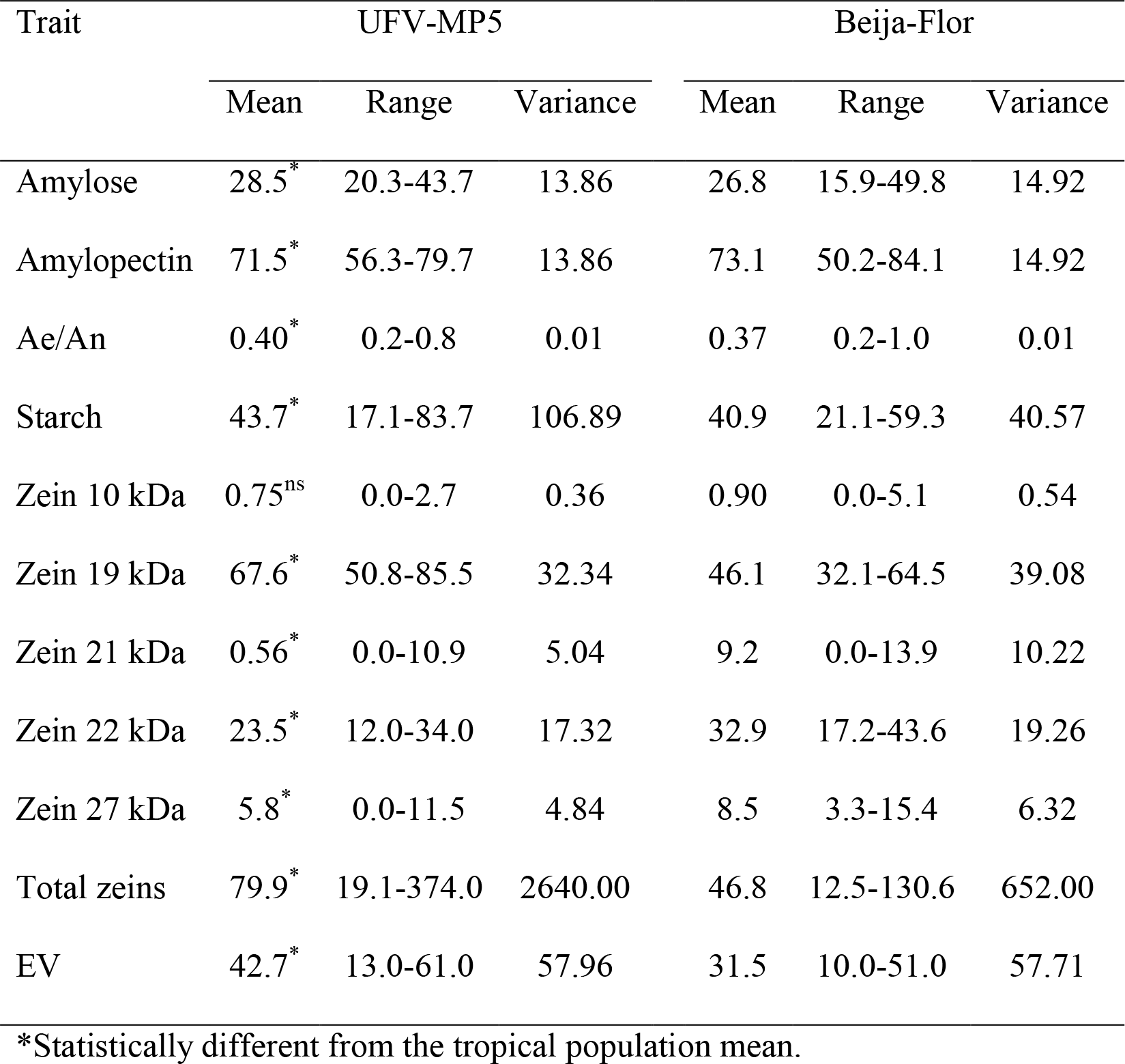
Average contents of amylose (%), amylopectin (%), starch (%), zein subunits (%), and total zeins (ng/mg of endosperm), amylose/amylopectin ratio (Ae/An), EV (mL/g), range (minimum-maximum), and phenotypic variances in the temperate and tropical populations

Significant associations were observed in the temperate population for the contents of 19 and 22 kDa zeins, starch, and amylose. Regarding the zein subunits, 102 significant associations were identified for the 19 kDa zein and 80 significant associations were identified for the 22 kDa zein, practically all located on chromosome 4. Sixty-eight of the significant associations were common to the two subunits. For the 19 kDa subunit, two associations were located on chromosome 1. For the 22 kDa subunit, one association was located on chromosome 1. Based on the significant associations, seven candidate genes related to the contents of these two subunits were identified. A candidate gene for total zein content was also identified. Eight significant associations were identified on chromosomes 1, 2, 3, 6, 8, and 9 for starch content. For amylose content, 19 significant associations were identified in all chromosomes. Two candidate genes for starch content and two for amylose content were found (Table 2). The candidate genes identified in the temperate population are associated with four biological processes, four molecular functions, and one cellular component. In addition, the proteins associated with these genes were classified into five categories, with a higher proportion of chaperones and protein-modifying enzymes (Online Resource Figure 3).

**Table 2.** SNP chromosome and position and candidate genes associated with starch, amylose, zein subunits, and total zeins.

| Population | Trait | Chr. | Position | Candidate Gene | Annotation |
| --- | --- | --- | --- | --- | --- |
| UFV-MP5 | Starch | 1 | 33454328 | Zm00001d028397 | Acyl-CoA-binding domain-containing protein 3 |
|  |  | 2 | 204459673 | Zm00001d004962 | malate dehydrogenase |
|  | Amylose | 6 | 158964536 | Zm00001d038512 | UDP-glycosyltransferase 85A7 |
|  |  | 10 | 138643984 | Zm00001d026113 | naked endosperm 2 |
|  | Total zeins | 4 | 828609 | Zm00001d048591 | chaperone dnaJ 6 |
|  | 19/22 kDa | 4 | 1457058 | Zm00001d048617 | ubiquitin transferase E3 type *RING* |
|  |  | 4 | 3199492 | Zm00001d048687 | *Heat Shock* 70 kDa |
|  |  | 4 | 6012358 | Zm00001d048812 | alpha-zein Z1_C1_7 |
|  |  | 4 | 6012386 | Zm00001d048813 | alpha-zein Z1_C1_8 |
|  |  | 4 | 6012386 | Zm00001d048817 | alpha-zein Z1_C1_19 |
|  |  | 4 | 7231847 | Zm00001d048894 | transcription factor MAD-BOX |
| Beija-Flor | Amylose | 10 | 26544422 | Zm00001d023892 | shikimate dehydrogenase1 |
|  |  | 10 | 73708417 | Zm00001d024486 | terpene synthase10 |
|  | Zein 21 kDa | 7 | 4201475 | Zm00001d018751 | amino acid permease 6 |

In the tropical population, there were six significant associations for the content of the 21 kDa subunit on chromosomes 5 and 7. Based on the physical position of the significant SNPs, a candidate gene associated with zein biosynthesis was identified (Table 2). Two significant associations were identified for amylose content on chromosome 10, providing two candidate genes.

## Discussion

There are several methods for quantifying zeins, starch, amylose and amylopectin. SDS-PAGE (Hamaker et al. 1995; Parsons et al. 2020), antibodies (ELISA method) (Wallace et al. 1990), reversed-phase liquid chromatography (RE-HPLC) (Flint-Garcia et al. 2009; Shawa et al. 2021), capillary electrophoresis (CE) (Parris et al. 1997), and mass spectrometry (MS) (Postu et al. 2019) have been used to characterize the distribution profile of zein subunits and determine zein contents. Regarding the quantification of starch, amylose and amylopectin, the iodometric method is still the main procedure in use (Bates et al. 1943; Morrison and Laignelet 1983; Sene et al. 1997; Zhu et al. 2008). The other includes the enzymatic method (Gibson et al. 1997), high-performance molecular size liquid chromatography (HPSEC) (Grant et al. 2002; Kobayashi et al. 1985), differential scanning calorimetry (DSC) (Sievert and Holm 1993), and near infrared spectroscopy (NIR) (Sampaio et al. 2018). In this study, microfluidic electrophoresis and Megazyme’s commercial Amylose/Amylopectin Kit, following the standard protocol, proved to be applicable to breeding programs, taking into account cost, practicality, and precision. The costs of and time for quantification, compared to measuring EV, limit quantifications to a scale of a few hundred selection units. Taking into account only the cost of acquiring the protein chips (250 samples) and the Megazyme kit (100 samples), the cost for zein quantification was 50% lower, approximately two dollars/sample. In terms of practicality, the average time spent characterizing ten accessions via microfluidic electrophoresis was four hours. The characterization performed by SDS-PAGE took approximately 72 h. The average time to quantify starch, amylose, and amylopectin for 14 plants was 8 h.

The characterization of a temperate and a tropical population revealed that across more than 100 years of successful breeding in the USA (Ziegler and Ashman 1994), based mainly on EV, indirect selection by popcorn breeders has increased the 19 kDa zein and total zein contents and decreased the fractions of other zeins, with slight changes in the starch and amylose/amylopectin contents. Based on molecular weight, the 19 and 22 kDa subunits were classified as alpha-zeins, and the 27 and 10 kDa subunits were classified as a gamma- and delta-zein, respectively (Shewry and Casey 1999). Among the zein subunits, alpha-zeins are associated with endosperm hardness and their content has a high correlation (0.96) with EV (Borras et al. 2006). Taking into account that starch is the major polymer involved in popcorn expansion (Sweley et al. 2013), popcorn breeding should be focused in increasing the 19 kDa alpha-zein, total zeins, and starch contents. North American and Argentine commercial hybrids have an average starch content of 64% and an amylose/amylopectin ratio of 27.5/72.5 (0.40) (Borras et al. 2006; Sweley et al. 2013), which is the average value observed in the temperate and tropical popcorn populations. The amylose/amylopectin ratio affects the quality and development of the endosperm (Li et al. 2018b), as well as properties useful to the food industry, such as gelatinization and solubilization, which are relevant aspects during the development of starch-based products (Lemos et al. 2019). In popcorn, the amylose content ranges from 27 to 45% (Borras et al. 2006; Park et al. 2000).

Alpha-zeins (19 and 22 kDa) are encoded by four gene subfamilies (Z1A, Z1B, Z1C, and Z1D) (Holding 2014), distributed in seven chromosomes (Li and Song 2020). The Z1A, Z1B, and Z1D subfamilies participate during the synthesis of the 19 kDa alpha-zein, and Z1C encodes the 22 kDa alpha-zein (Holding 2014). The 19 kDa alpha-zein is related to 25 genes in five genomic regions distributed on chromosomes 1, 4 and 7, with an approximate coverage of one Mb (Song and Messing 2002). Copies of genes from the Z1A and Z1C subfamilies are in two positions on chromosome 4 (Miclaus et al. 2011). The candidate genes Zm00001d048812 (Z1_C1_7), Zm00001d048813 (Z1_C1_8), and Zm00001d048817 (Z1_C1_19) are described as belonging to the Z1C subfamily and are directly related to the synthesis of alpha-zeins. The Z1_C1_8 gene is also involved in the biosynthesis of the 22 kDa alpha-zein (Song et al. 2001).

The Z1C subfamily consists of 16 genes. Of these, six are expressed during endosperm development (azs22.4, azs22.7, azs22.8, azs22.9, azs22.19 and fl2-azs22.16) (Feng et al. 2009; Song and Messing 2003). From the gene expression analysis in immature embryos for the *locus* Z1C1, it was demonstrated that the genes azs22.4 and azs22.19 (Z1_C1_19) had the highest levels of transcription (Miclaus et al. 2011). Although it has been demonstrated that the Z1A *locus* presents a higher level of transcription and have a major role in the synthesis of alpha-zeins, whose synthesis is dependent on the joint action of other *loci*, Z1_C1_19 is a key gene in the process of alpha-zein synthesis. The azs22.8 gene accounts for approximately 13% of the genes belonging to the Z1C subfamily (Feng et al. 2009). The expression of this gene peaks at 18 days after pollination, concomitant with the peak expression of the fl2 gene (Feng et al. 2009), suggesting the joint action of these genes in the synthesis of alpha-zeins. In addition, Segal et al. (2003) silenced this gene via RNA interference, promoting a reduction in the expression of the 22 kDa alpha-zein. Therefore, it could be concluded that this gene is directly related to alpha-zeins synthesis. The azs22.7 gene accounts for 12.6% of the genes belonging to the Z1C subfamily and its peak expression occurs 22 days after pollination. Furthermore, it seems to act together with the azs22.19 gene during alpha-zein biosynthesis (Feng et al. 2009).

The candidate gene Zm00001d048687 encodes the 70 kDa heat shock protein (HSP70). This protein belongs to the HSP70 family, is described as a chaperone (Boston et al. 1991) and acts on the processing of proteins in the endoplasmic reticulum. The overexpression of HSP70 was observed in mutants with low zein content (*fl2, De-B30*, and *Mc*), suggesting a role of this protein in the transport and/or accumulation of zeins in protein bodies. An increase in the synthesis of chaperones and a reduction in the biosynthesis of zeins were also observed in other mutants, such as *Opaque1* and *Opaque10* (Shewry and Casey 1999), suggesting that there is a relationship between this family of proteins and the regulation of zein biosynthesis. Thus, the DNAJ6 chaperone protein encoded by the Zm00001d048591 gene may also be related to zein biosynthesis, as chaperones contribute to the folding and deposition of zeins in protein bodies (Müntz 1998).

In addition to the genes involved in zein synthesis, four types of transcription factors are known to act directly in the transcriptional regulation of zein genes (*O2, PBF1, OHP1/2, and MADS-box*) (Li et al. 2018a). The transcription factor zmMADS47, a member of the MADS family, controls the activation of genes that encode the synthesis of alpha-zeins and the 50 kDa zein and interacts with the *O2* mutant (Li and Song 2020; VicenteCarbajosa et al. 1997). The candidate gene Zm00001d048894 encodes a transcription factor of the MAD-BOX family, located in the endosperm, and has protein dimerization activity, acting in the development of the endosperm. Zm00001d048617 encodes a protein in the *RING zinc finger domain* superfamily. These proteins play crucial roles in plant responses to drought stress (Kong et al. 2013). VicenteCarbajosa et al. (1997) demonstrated that a protein in this superfamily binds to Prolamin-BOX, a region rich in promoters of genes related to seed storage proteins in maize, and interacts with transcriptional activators of *Opaque2*. This suggests that other transcription factors in addition to *Opaque2* may coordinate the expression of zein genes, such as those identified.

A candidate gene was identified for the 21 kDa subunit. Zm00001d018751 codes for the protein permease of amino acid 6 (Aap6), a member of the amino acid transporters family (ATFs) (Ortiz-Lopez et al. 2000). Peng et al. (2014) demonstrated that Aap6 acts as an important regulator in the synthesis and accumulation of storage proteins in rice grains as well as in starch biosynthesis. According to Yao et al. (2020), ATFs exhibit biochemical similarities in different species (e.g., rice, arabidopsis, and soybean), suggesting that the function of these proteins is conserved among vascular plants. Thus, by homology, it is assumed that Aap6 is related to the biosynthesis and accumulation of zeins.

Concerning starch content, candidate genes Zm00001d028397 and Zm00001d004962 were identified. Zm00001d028397 encodes the Acyl-CoA-3 protein (ACBD3). In maize, acyl-CoA binding proteins are encoded by nine genes distributed in five chromosomes, with three genes located on chromosome 1, including ACBD3 (Okita 1992; Sung et al. 1988; Zhu et al. 2021). Although there are no studies involving the association of this protein with starch content in maize, Zhu et al. (2021) observed high expression of this protein during endosperm development (18 to 22 days after pollination) and associations with grain-related traits, such as weight, length and width, number of seeds per row and oil content, suggesting that this protein has multiple functions during seed development. Zm00001d004962 encodes the enzyme malate dehydrogenase, which acts in the gluconeogenesis pathway. This biosynthetic pathway is responsible for the generation of glucose from non-sugar carbon substrates, such as pyruvate, (s)-lactate, and glycerol. In turn, glucose acts in the synthesis of sucrose, which plays the role of substrate for storage glycosides such as starch and fructans (Okita 1992; Sung et al. 1988). Thus, this enzyme is indirectly related to starch biosynthesis.

The candidate genes Zm00001d038512, Zm00001d026113, Zm00001d023892 and Zm00001d024486 are related to amylose content. Zm00001d038512 encodes the protein o-cytokine glycosyltransferase, a member of the UDP-glycosyltransferase family. Glycosyltransferases catalyze the formation of glycosidic bonds, generating disaccharides, oligosaccharides and polysaccharides (Lairson et al. 2008). In a study by Li et al. (2018b), four of the 39 candidate genes related to amylose content encode glycosyltransferases. This class of proteins plays an important role in amylose metabolism, acting together with granule bound starch synthase (GBSS), the main enzyme responsible for the synthesis of amylose (Li et al. 2018b). The transcription factor NKD2 (Naked Endosperm 2), encoded by the candidate gene Zm00001d026113, plays a key role in endosperm development, including the deposition of storage nutrients (Gontarek et al. 2016). Gontarek et al. (2016) observed a reduction in the expression of genes related to storage nutrients in maize grains, for example, GBSS and prolamin-box-binding transcription factor, in NKD1/NKD2 mutants. Consequently, there was a 32.5% reduction in the starch content of the endosperm.

The candidate gene Zm00001d023892 encodes the enzyme shikimate dehydrogenase, which catalyzes the fourth reaction of the shikimic acid pathway (Fonseca et al. 2006). This pathway is the main metabolic route for the synthesis of phenolic compounds, linking the carbohydrate precursors derived from glycolysis to the biosynthesis of aromatic compounds (Parish and Stoker 2002). The shikimate pathway begins with two glucose metabolites, phosphoenolpyruvate (PEP) + D-erythrose-4-phosphate (E4P), and at the end of the pathway, chorismic acid (CHA), a precursor molecule of different metabolites, is obtained (Bentley 1990; Ducati et al. 2007; Parish and Stoker 2002). Among these, phenylalanine stands out playing a role in the biosynthesis of acetyl-CoA, a molecule originating from carbohydrate metabolism (Coggins et al. 2003; Ducati et al. 2007).

In the terpene biosynthetic pathway, initiated by the condensation of acetyl-CoA molecules, the terpene synthase enzyme, encoded by the candidate gene Zm00001d024486, acts in the conversion of intermediate molecules into dimethylallyl pyrophosphate and isopentenyl pyrophosphate (Pyne et al. 2019). These terpenes, in turn, are precursors in the biosynthesis of molecules that act as glucosyl transporters in the process of N-glycosylation, the reaction of adding carbohydrates to protein and lipid molecules (Guimaraes et al. 2019; Tholl et al. 2006).

Concluding, microfluidic electrophoresis and the commercial Amylose/Amylopectin Kit developed by Megazyme can be used in popcorn breeding, taking into account cost, practicality, and precision. Notably, the difference in quality between temperate and tropical popcorn populations is due in part, by indirect selection based on EV, to an increase in the 19 kDa alpha-zein content and total zeins content and a reduction in other zein fractions, without significant changes in starch and amylose/amylopectin contents.

## Acknowledgements

I thank the National Council for Scientific and Technological Development (CNPq), the Brazilian Federal Agency for Support and Evaluation of Graduate Education (Capes; Finance Code 001), and the Foundation for Research Support of Minas Gerais State (Fapemig) for financial support.

## Competing Interests

The authors have no relevant financial or non-financial interests to disclose.

## Author contributions

LFG: methodology, analysis, and writing - original draft preparation; JMSV: conceptualization, funding acquisition, and writing - review and editing; MPR: metodology; RBC: methodology; HJOR: methodology and writing - review and editing; JLRF: methodology and writing - review and editing. All authors read and approved the final manuscript.

Data Availability The dataset is available at https://doi.org/10.6084/m9.figshare.8277629.v2.

---

**Suplementary Figure 1.**
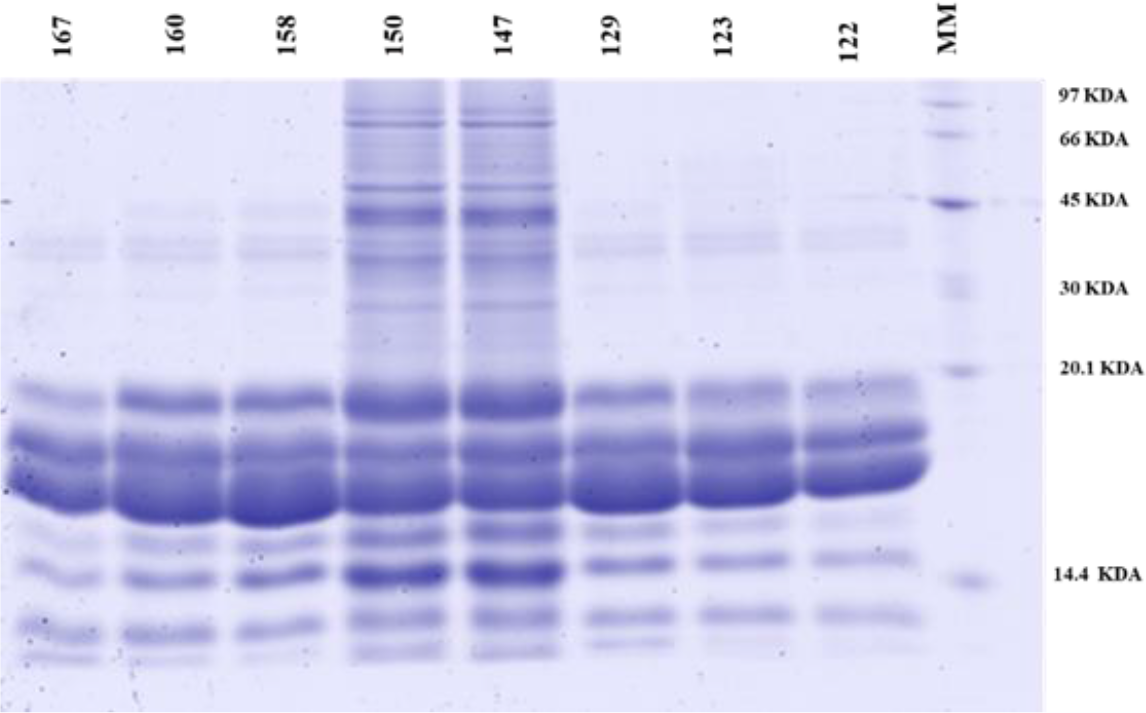
Zein profile on SDS-PAGE gel of eight individuals from the temperate population and molecular marker (MM) GE Healthcare.

**Suplementary Figure 2.**
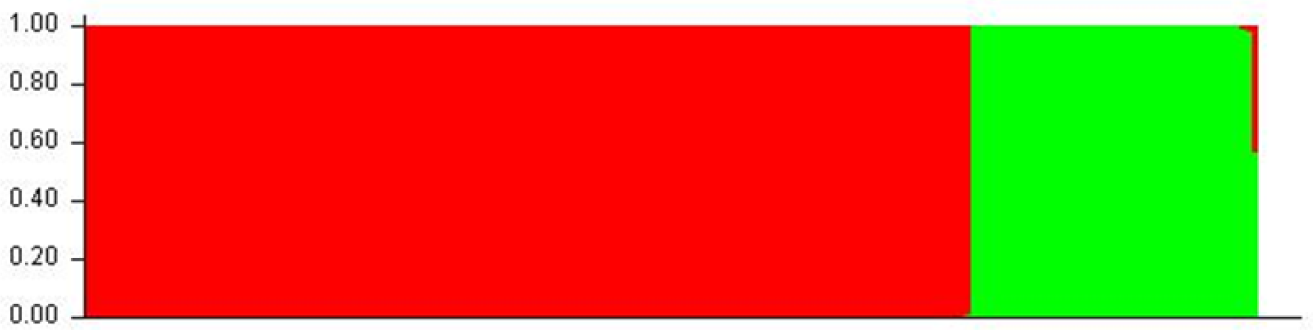
Subpopulation structure identified in the temperate population.

**Suplementary Figure 3.**
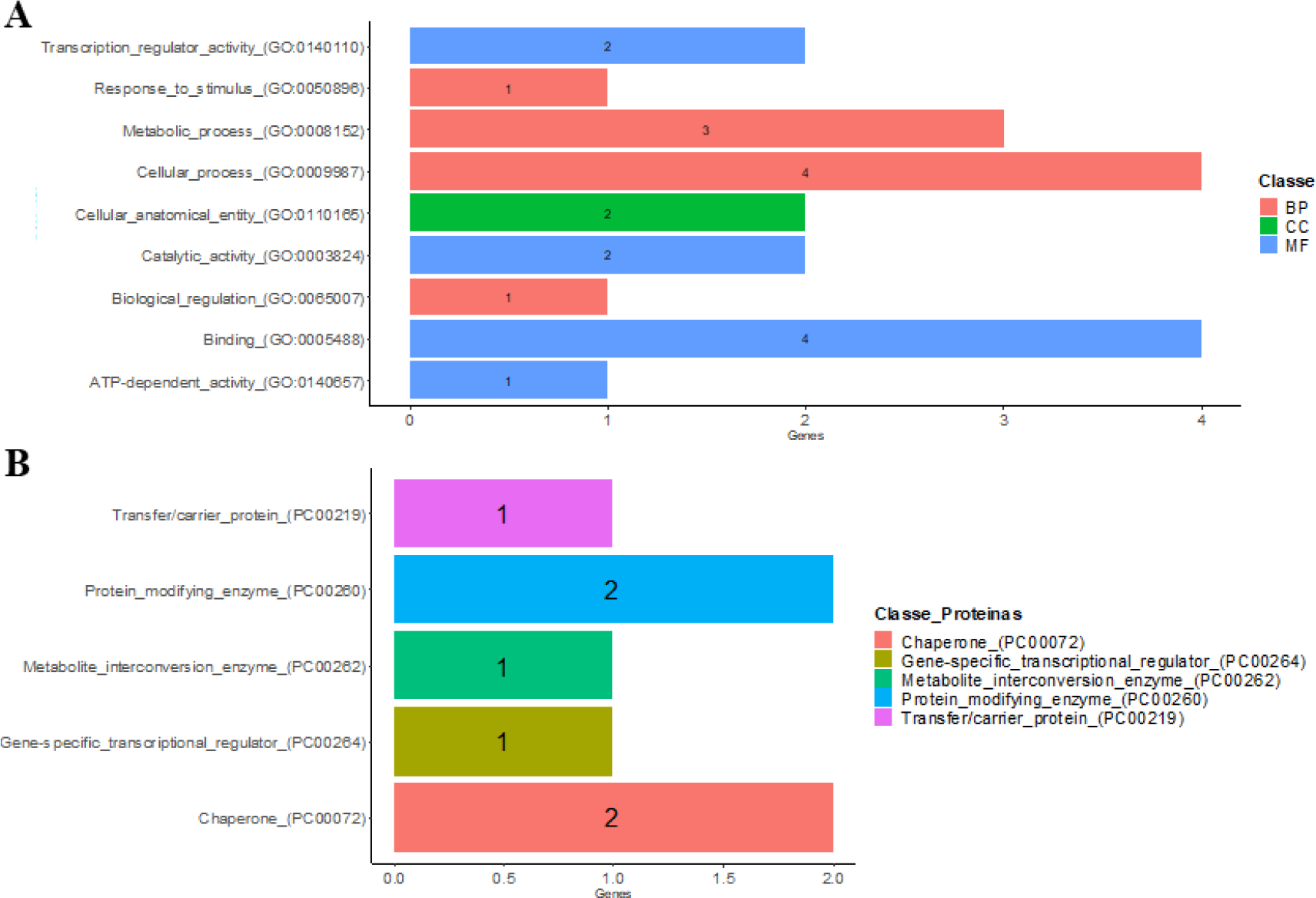
Candidate genes ontology.

## Notes

https://doi.org/10.6084/m9.figshare.8277629.v2

### Competing Interest Statement

The authors have declared no competing interest.

